# A linear programming-based strategy to save pipette tips in automated DNA assembly

**DOI:** 10.1101/2021.10.13.464196

**Authors:** Kirill Sechkar, Zoltan A. Tuza, Guy-Bart Stan

## Abstract

Laboratory automation and mathematical optimisation are key to improving the efficiency of synthetic biology research. While there are algorithms optimising the construct designs and synthesis strategies for DNA assembly, the optimisation of how DNA assembly reaction mixes are prepared remains largely unexplored. Here, we focus on reducing the pipette tip consumption of a liquid-handling robot as it delivers DNA parts across a multi-well plate where several constructs are being assembled in parallel. We propose a linear programming formulation of this problem based on the capacitated vehicle routing problem, along with an algorithm which applies a linear programming solver to our formulation, hence providing a strategy to prepare a given set of DNA assembly mixes using fewer pipette tips. The algorithm performed well in randomly generated and real-life scenarios concerning several modular DNA assembly standards, proving capable of reducing the pipette tip consumption by up to 61% in large-scale cases. Combining automatic process optimisation and robotic liquid-handling, our strategy promises to greatly improve the efficiency of DNA assembly, either used alone or in combination with other algorithmic methods.

## I. INTRODUCTION

Among the key factors driving synthetic biology research, which is concerned with efficiently engineering organisms with useful new properties, is the automation of laboratory processes. Allowing to produce much larger and higher-quality experimental datasets, it not only accelerates the rate of research, but also enables the use of computational analysis tools that require large amounts of experimental data, which is difficult to produce in a reasonable time by low-throughput human manual labor[1].

Implementing a genetic circuit in living organisms typically involves DNA assembly – the joining of DNA fragments (“parts”) where each part is usually a functional genetic element, into a longer nucleic acid chain (‘construct’), which can then be introduced into a host organism to induce the desired phenotype. Likewise to other aspects of synthetic biology research, DNA assembly has been greatly impacted by the adoption of laboratory automation practices – namely, the use of liquid-handling robotics [2]. A particular advantage of using such robots is the parallel preparation of multiple different constructs in distinct wells of a single multi-well array plate, which greatly improves the throughput and time efficiency of DNA assembly.

Metrics quantifying the cost and time benefits of executing a given DNA assembly scenario by a liquid-handling robot instead of a human worker have been proposed previously [3]. Furthermore, recent years have seen the rise of algorithms aiming to improve the efficiency of automated DNA assembly. Primarily, they deal with the design stage of the research cycle: e.g., the number of reactions required to obtain the desired construct in a multi-step assembly can be minimised by representing possible sequences of assembly steps as graphs and selecting the optimal one [4]. Alternatively, optimisation occurs on the high level of the design implementation phase. For instance, when testing different combinations of DNA parts to identify the most successful design, the most promising constructs worth assembling and testing in practice can be identified *a priori*, so that there is no need to implement inauspicious designs [3]. Meanwhile, lower-level optimisation of a liquid-handling robot’s exact actions as it sets up reaction mixes for parallel DNA assembly has largely remained neglected. This contrasts with the general practice of process automation, where compilers optimize the commands and execution order of human-written programs on several levels at once [5].

Many of the widely adopted DNA assembly standards (e.g., BASIC [2], MoClo [6] or Start-Stop assembly [7]) are “one-pot”, i.e. involve mixing all of a construct’s DNA parts together before ligating them in a single reaction. Therefore, setting up parallel assembly reactions on a well plate involves preparing different mixtures of several DNA parts in different wells. Although during a one-pot assembly a single part is most often required in multiple distinct constructs, reusing the same pipette tip to deliver a part to several wells introduces a high risk of contaminating a master mix by the DNA parts present in the previously visited wells. Therefore, either a new tip is used for delivery to each well, which results in an excessively large uptake of pipette tips and thus increased operating costs, or the pipette is washed after every delivery, which can be time-consuming.

The need for washing or changing the tip after each DNA part addition can be avoided. Suppose that at some point in the robot’s program a single tip has distributed a certain DNA part to several wells and now proceeds to the next well. If none of the previously visited wells contain any DNA parts that have not already been added to the next well earlier in the laboratory robot’s program, there is no risk of contaminating the next well by any DNA part that would not be present in it anyway. Fig. 1 showcases the minimisation of pipette tip consumption for a simple example of parallel DNA assembly of two two-part constructs following this principle.

**Fig. 1.**
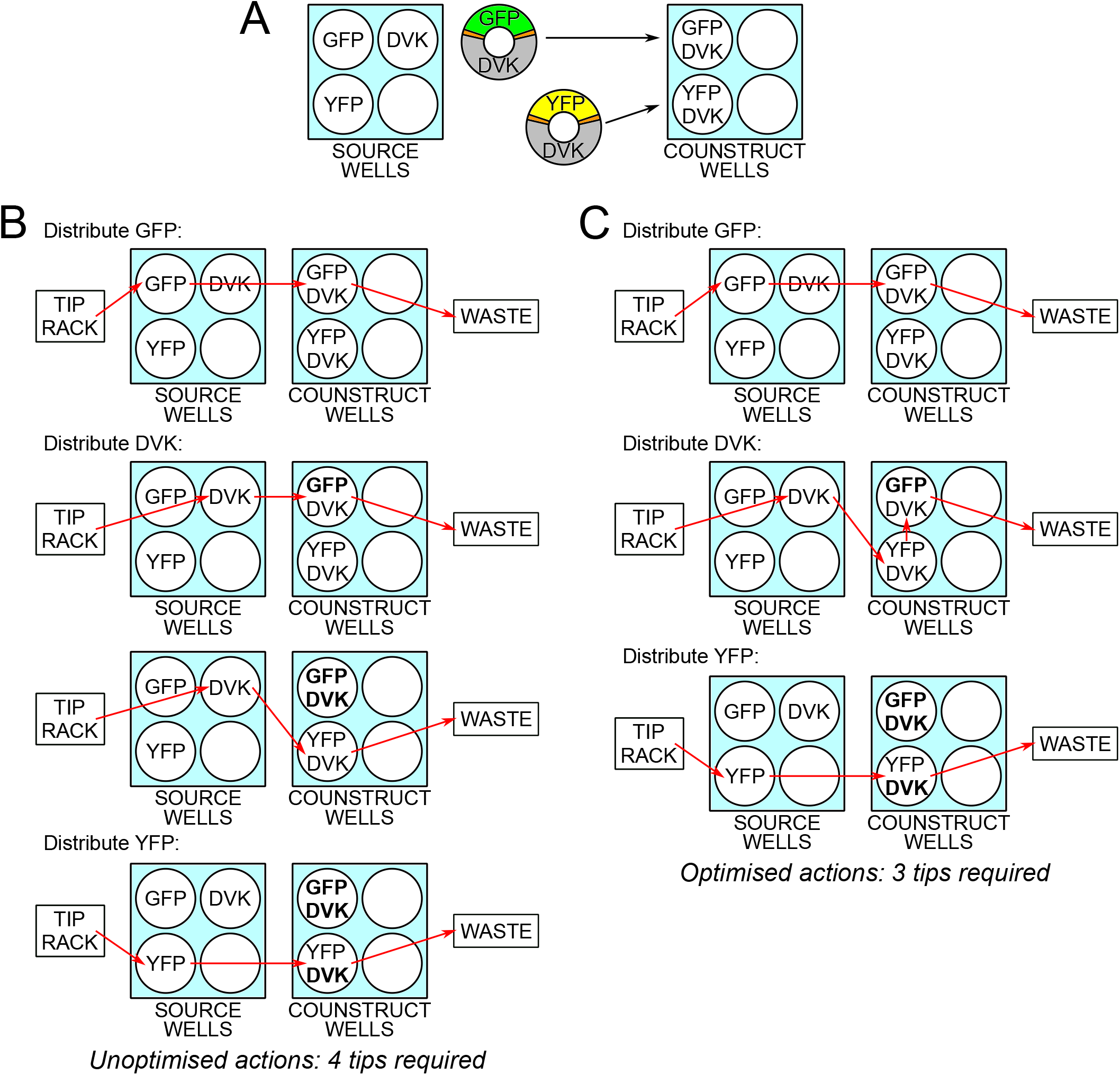
An instance of pipette tip consumption minimisation. a) One construct comprises the Green Fluorescent Protein (GFP) gene inserted into the DVK vector backbone, while the other has the same backbone hosting the Yellow Fluorescent Protein (YFP) instead. b) Unoptimized sequence of DNA part additions. Source wells display which DNA part solution they contain; construct wells display which DNA parts must be added to them. The parts already present in the wells at the time of a given part’s addition are displayed in bold. The route of each pipette tip is outlined by arrows starting at the tip rack, then collecting the relevant DNA part solution from a source well, dispensing it to the construct wells, and finally being discarded into the waste. c) If the series of DVK additions is optimized, three pipette tips can be used instead of four without the risk of contamination.

Therefore, for every DNA part, the order in which it is delivered to the construct where it is required in can be optimized so as to minimize the number of pipette tips needed to perform all of its additions. This calls for an algorithm that receives a description which DNA parts have to be added to which construct wells, and produces a sequence of pipette commands such that for each DNA part involved the order of actions is optimized, thereby minimising the pipette tip consumption of the liquid-handling robot. In this paper, we propose such an algorithm based on the capacitated vehicle routing problem (CVRP) [8].

## II. METHODS

### A. The capacitated vehicle routing problem (CVRP)

Let us define the CVRP before we show how its LP formulation can be modified to describe the pipette tip-saving problem. Essentially, the vehicle routing problem can be described as follows. There is a *depot*, where *goods* are stored, and *customers*, to whom the goods must be delivered. The customers are connected among themselves and to the depot by *roads*, each of which has a cost assigned to it. The objective is to use a fleet of *vehicles*, which can travel the roads, to deliver the goods to all customers while minimising the sum of the costs of all roads that are travelled by the vehicles, which is the problem’s objective function. The constraints are that the number of vehicles in the fleet is fixed and that every vehicle must start and end its journey at the depot. In the capacitated vehicle routing problem, it is additionally specified that each vehicle can only carry an amount of goods that does not exceed the *vehicle capacity*.

The road network can be represented as a graph *G* = (*V, E*), where customers are nodes {*v*_1_, *v*_2_ … *v*_*n*_} ∈ *V* with the depot indexed as the node *v*_0_ *V* ∈ The roads between them are edges {*e*_*ij*_ | 0 ≤*i, j ≤n*} ∈ *E*, where *e*_*ij*_ is the edge from node *v*_*i*_ to node *v*_*j*_ (note that *e*_*ij*_ and *e*_*ji*_ are two distinct edges) and the cost of the edge *e*_*ij*_ is given by the number *cost*(*e*_*ij*_) = *c*_*ij*_. The fleet size can be expressed as *K* and the vehicle capacity as *κ*. Finally, let us have variables {*x*_*ij*_ | 0 ≤*i, j ≤n*}, where *x*_*ij*_ = 1 if the edge *e*_*ij*_ is travelled by any vehicle in the fleet and *x*_*ij*_ = 0 otherwise. Then, the objective of the CVRP is to determine the values of {*x*_*ij*_} that minimize *C* = ∑*x*_*ij*_*c*_*ij*_ subject to linear constraints defined on *x*_*ij*_, *K* and *κ* [8].

### B. Casting the pipette tip-saving problem as a modified CVRP

The problem of pipette tip-efficient delivery of DNA parts to the wells that need them can be cast as a slightly modified CVRP.

Let us consider a numbered list outlining the order in which the parts are delivered. For the part number *h* in this list, let us construct a graph *G*_*h*_ = (*V, E*) analogous to that of the CVRP. An aliquot of the liquid solution containing the DNA part being distributed corresponds to a “good” that needs to be delivered to a “customer” node *v*_*i*_, where the “customers” correspond to the wells where an assembly mix is prepared. Meanwhile, every pipette tip used corresponds to a “vehicle”, delivering the DNA part solution (“goods”) to the construct wells (“customers”). As any pipette tip must start its journey at the DNA part solution source and end it at the waste bin, let us denote both these destinations as a single “depot” node *v*_0_, from which each “vehicle” trajectory must originate and where it must also end. As a single pipette tip’s volume is finite, there is a maximum number of wells it can serve, which defines its capacity *κ*.

Finally, the edges correspond to pipette transitions between wells, while their costs describe potential contamination events. Contamination may arise when the same pipette tip first visits a well containing a certain DNA part (one of the *h* − 1 DNA parts already added to the mixes requiring them) and then goes to a new well which does not have it, hence the latter well being contaminated by this part. More formally, for every well *v*_*a*_ we can write down a set 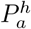 listing all DNA parts present by the time the robot proceeds to distribute DNA part number *h*. If well *v*_*i*_ has been visited by the same tip before well *v*_*j*_ is visited, then *v*_*j*_ would be contaminated if 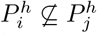. To keep track of all such contaminations, let there be an edge *e*_*ij*_ ∈ *E* between any two wells *v*_*i*_, *v*_*j*_ *∈ V*, such that *cost*(*e*_*ij*_) = 1 if 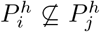, and *cost*(*e*_*ij*_) = 0 otherwise (any edge from or to the “depot” – *e*_0*i*_ or *e*_*i*0_, respectively – has cost 0 by definition, since collecting the part solution from the source or trashing the tip is irrelevantto well crosscontamination). As long as a single tip’s route includes only zero-cost edges, then for any well 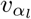 we know that none of the previously visited wells 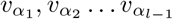 contain DNA parts not present in 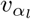 already: indeed, since 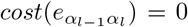, we have 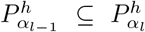; then, since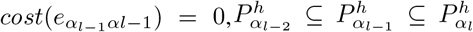. We can proceed by induction until we obtain 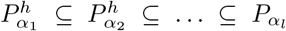, which excludes the contamination condition.

The objective of the pipette tip-saving problem is to find the variables {*x*_*ij*_} (where, as previously, *x*_*ij*_ indicates whether the edge *e*_*ij*_ is travelled by any of the “vehicles”) such that the number of tips used is minimized while cross-contamination is avoided and no tip serves more wells than its capacity *κ* allows. This means that our problem’s formulation is identical to the CVRP, except for two changes. First, the total cost of all the travelled edges *C* is fixed at zero to avoid contamination; second, the “fleet size” (number of pipette tips used) *K* is not a fixed value, but the objective function to be minimized.

### C. Solving the pipette tip-saving problem as a Linear Program

The advantage of expressing our pipette tip-saving problem as a Linear Program (LP) is that LPs constitute a well-studied class of problems, for which numerous powerful numerical solvers have been developed. Therefore, the LP formulation of the problem of distributing a single DNA part to the construct wells can be leveraged to formulate an algorithm that makes use of an LP solver to minimise pipette tip consumption over the whole procedure of preparing DNA assembly master mixes. We implemented such an algorithm in Python, using Gurobi™ Optimizer’s LP solver [9].

Briefly, the algorithm is provided with the composition of the DNA constructs to be assembled, as well as the location of the wells where the corresponding mixes should be prepared and of the sources of DNA part solutions. From this input, it determines a list describing which DNA part will be distributed first across all the wells that require it, which part will be delivered second, and so on. Now, for every part in this list, an LP problem is solved, which allows to determine the best way of delivering it. Reaching the end of the part list means that we have obtained a sequence of liquid-handler commands, enabling the contamination-free distribution of all parts and reducing the number of pipette tips used.

The algorithm is described in more detail in Fig. 2, while Fig. 3 demonstrates our algorithm in action for the example considered in the introduction.

**Fig. 2.**
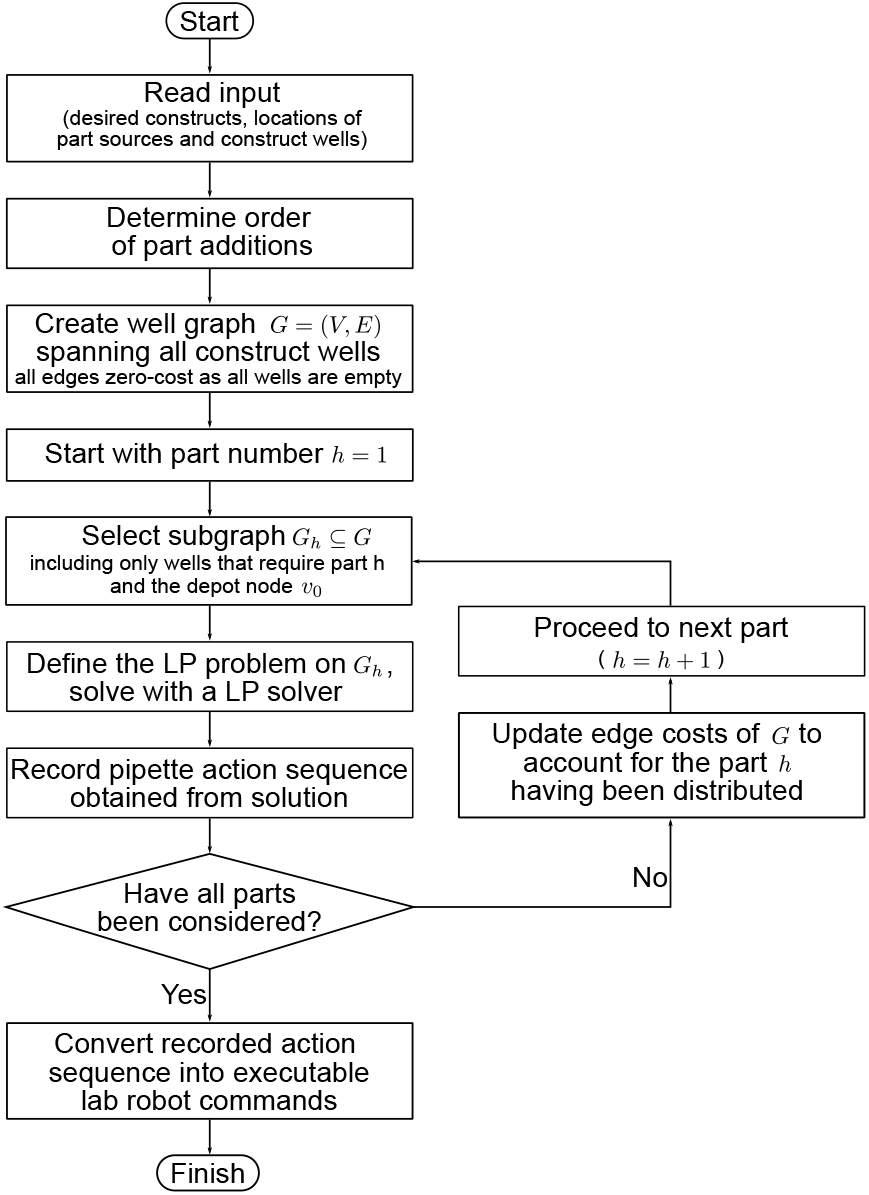
Flowchart describing our proposed pipette tip consumption minimization algorithm.

**Fig. 3.**
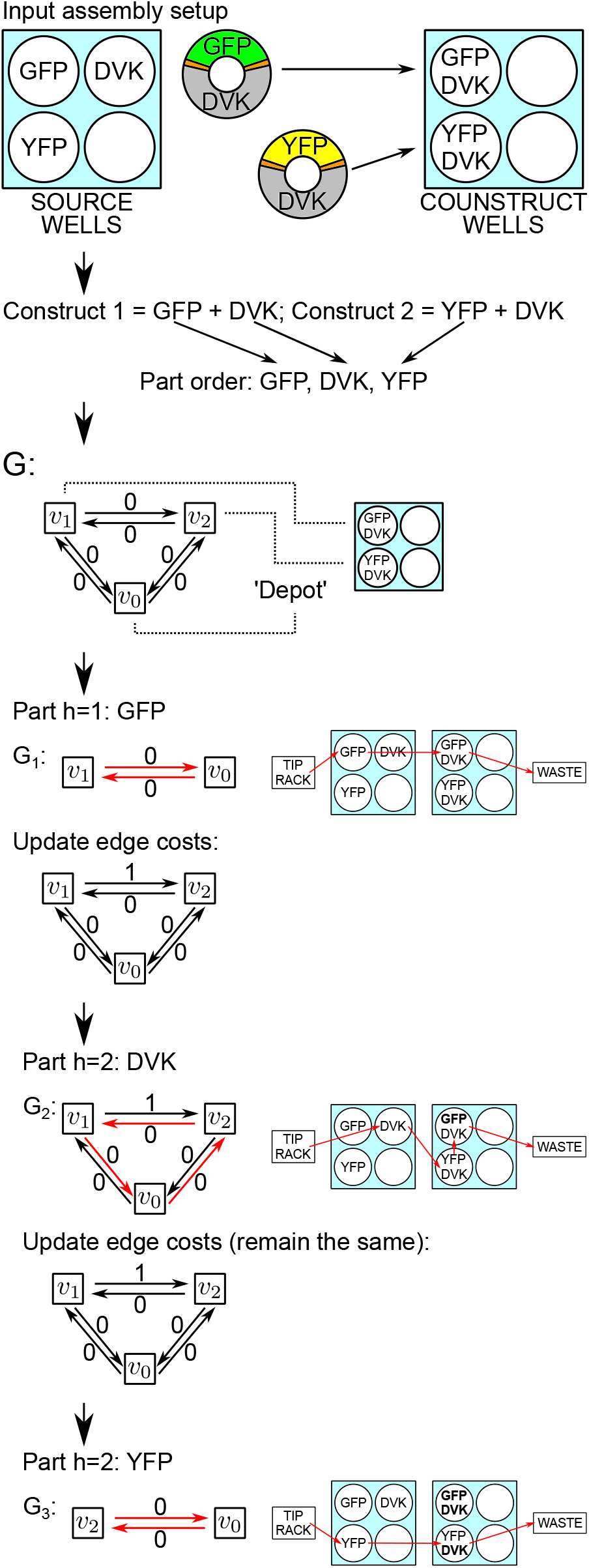
The algorithm applied to an example assembly setup – in the end, the optimised series of pipette commands, which uses only 3 tips to perform 4 part additions is produced (same solution as in Fig. 1). The edges travelled by a pipette tip in the LP problem’s solution are displayed in red. Note how the cost-one edge is avoided in *G*_2_, thereby preventing contamination.

### D. Testing the pipette tip-saving algorithm’s performance

To evaluate the performance of our algorithm, we implemented it in Python and tested it on randomly generated inputs for the Start-Stop DNA assembly protocol [7]. Each assembled gene we considered in our tests consists of four parts: a promoter, a ribosome-binding site (RBS), a coding sequence (CDS) and a terminator. In this example, a construct is generated by picking one of 6 possible promoters, one of 4 RBSs, one of 3 CDSs and one of 3 terminators, reflecting the sizes of DNA part libraries used in a past experiment in our labs. For every number of constructs from 2 to 96, we generated 50 random assembly scenarios with this many random constructs assembled in parallel. For each of these sets of 50 optimisations, the mean percentage of pipette tips saved relative to the case where each part addition is performed with a fresh tip was calculated and plotted in Fig. 4.

**Fig. 4.**
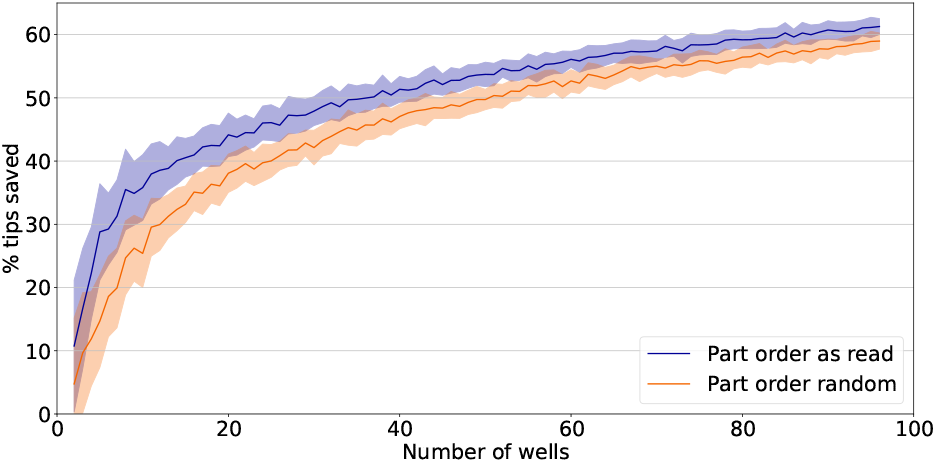
Results of performance testing with randomly generated Start-Stop assembly inputs, for 2 up to 96 paralelly assembled constructs. The shading shows the range of the percentages lying within the standard deviation from the mean.

To demonstrate that the algorithm can save tips in real cases as well as for randomly generated ones, we turned to the packages DNA-BOT [2] and “OT2 Modular Cloning (MoClo) and Transformation in *E. coli* Workflow” [10], which automate the DNA assembly of the BASIC and MoClo standards respectively for the Opentron OT-2 liquid-handling platform. They contain test examples of known parallel DNA assembly scenarios, which can be used to assess the automation pipelines’ accuracy and efficiency – the complete sets of inputs can be found in these packages’ code repositories [2, 10]. The result of optimising the pipette movements for the situations described by them are displayed in Table I and Table II.

**TABLE I.** RESULTS OF PERFORMANCE TESTING WITH BASIC ASSEMBLY TEST INPUTS.

| Part Order | % tips saved |
| --- | --- |
|  | BASIC<br>5 parts<br>(max. 440 tips<sup>a</sup>) |
| as read | 49.54 |
| random | 50.68 |

<sup>a</sup>Pipette tips consumption when a fresh tip is used for each part addition.

**TABLE II.** RESULTS OF PERFORMANCE TESTING WITH MOCLO ASSEMBLY TEST INPUTS. THE EXAMPLE CONSTRUCTS WERE SPLIT BY THE NUMBERS OF PARTS THEY COMPRISE.

| Part Order | % tips saved |  |  |
| --- | --- | --- | --- |
|  | MoClo<br>2 parts<br>(max. 48 tips<sup>a</sup>) | MoClo<br>5 parts<br>(max. 120 tips<sup>a</sup>) | MoClo<br>8 parts<br>(max. 192 tips<sup>a</sup>) |
| as read | 31.25 | 50.00 | 54.69 |
| random | 27.1 | 28.3 | 38.2 |

<sup>a</sup>Pipette tips consumption when a fresh tip is used for each part addition.

To assess the effect of our method on determining the order of DNA part addition from the input data (from the part first encountered in the list of constructs to the last one), for every input the optimisation was also performed with a randomly permuted order of the input list. Hence, Fig. 4 and Table I each display two different sets of results, labelled “part order as read” or “part order randomly permuted”, respectively.

The results in this paper were produced on a server with 2x Intel^®^ Xeon^®^ CPU E5-2670 2.60GHz processors with 128 GB RAM, running on Ubuntu 18.04.3 LTS. The algorithm was implemented in Python 3.8. The Python packages numpy 1.17.2, scipy 1.6.0, tspy 0.1.1.1 and gurobipy 9.1.1 were used. When integrating the algorithms into the DNA-BOT and MoClo assembly pipelines, the packages opentrons 3.21.1, pyyaml 5.4.1 and pandas 0.25.1 were additionally used.

## III. RESULTS AND DISCUSSION

### A. Algorithm testing results

With more than half of pipette tips saved for the inputs describing high numbers of parallelly prepared Start-Stop assembly constructs, the results demonstrate the algorithm’s potential to significantly improve resource efficiency for parallel DNA assembly under various standards. As the number of constructs assembled in parallel grows, the tip savings rise – a likely explanation for this is that with higher input sizes, a single part is found in more constructs being prepared, and solving the CVRP-like problem on a larger subgraph can potentially yield longer chains of wells which the same pipette tip can visit. Therefore, the benefits of algorithmic pipette tip uptake optimisation could be expected to improve even more as synthetic biology research shifts towards larger-scale experiments and high-throughput screening [11, 12].

While even with DNA parts listed in random order considerable optimisation was achieved, using the part order as read from the input resulted in greater pipette tip savings in all but one scenario considered. Therefore, there may exist patterns of part order listing that on average increase pipette tip savings. In the future, identifying and exploiting such input patterns alongside our proposed LP-based optimisation of actions to distribute an individual part could yield an even more powerful pipette tip-saving method.

### B. Discussion

We propose an LP formulation of the problem of saving pipette tips while delivering a single part to wells where one-pot DNA assembly reactions are being performed in parallel. We provide performance test results for both randomly generated and real-life examples, demonstrating that algorithms leveraging this formulation in conjunction with a linear programming solver can be used to significantly reduce pipette tip consumption.

Underlying this potentially powerful DNA assembly optimisation strategy is the combination of a robotic liquid-handling platform and automatic process optimisation. Apart from being slower than a laboratory robot, a human worker with a conventional pipette can only collect a specified volume of solution and then dispense it all into the desired well, while distributing exact volume fractions across several destinations is typically not done manually. On the contrary, robotic liquid handler’s regulated pumps allow to perform this routinely. Consequently, having a robotic liquid handler follow simple pipetting protocols tailored for a human worker does not result in optimal procedure execution, leaving open avenues to realize the full potential of laboratory automation. Meanwhile, manually optimising the laboratory protocol instead of using automated algorithmic solutions would be a repetitive procedure required for every new assembly setup, becoming especially tedious and challenging for large-scale assemblies with dozens of potential contaminants to keep track of.

Our algorithm currently focuses on the lowest level of protocol optimisation, as it manages individual pipette actions for the distribution of a single part at a time. It is also not restricted to any single one-pot DNA assembly method. This allows to achieve optimisation of pipette tip consumption for a variety of different DNA assembly automation pipelines, as well as to combine our algorithms with higher-level optimisation strategies (which can determine which constructs should be synthesized and in which order [3, 4]) to achieve a truly multimodal optimisation of automated DNA assembly.

The proposed algorithm’s implementation in Python code is available via our open-access GitHub repository `pipette_opt` [13]. Our package offers API solutions which allow to incorporate our algorithm for pipette tip consumption minimization into existing DNA assembly pipelines for the Opentrons OT-2 liquid handler, namely DNA-BOT [2] (v1.0.0) and “OT2 Modular Cloning (MoClo) and Transformation in *E. coli* Workflow” [10]. Requiring only minor changes to the packages’ original code, `pipette_opt` allows the user to determine the optimized sequence of pipette actions during the DNA part distribution step, and translate it into Opentrons OT-2 commands. When the protocols generated by the modified pipelines were tested *in silico* using the Opentrons OT-2 simulation feature, we could indeed observe a significant reduction in pipette tip consumption.

## Supporting information

Start-Stop Assembly Random Inputs

Random Input Testing Results

## ACKNOWLEDGMENT

We thank Duncan Ingram for helping to prepare with the figures for this article, and Nicolas Kylilis and Eliza Atkinson for insightful discussions about DNA assembly. GBS gratefully acknowledges the support of the Royal Academy of Engineering through his Chair in Emerging Technologies (RAEng CiET 1819*\*5).

## Notes

### Competing Interest Statement

The authors have declared no competing interest.

