## Supporting Text for "A linear programming-based strategy to save pipette tips in automated DNA assembly"

#### Contents

**1 Linear Programming Problem Formulation** **S-1**

**2 Determination of the order of DNA part distribution** **S-4**

**3 Annotation to** `Start-Stop_Assembly_Random_Inputs.csv` **S-5**

**4 Annotation to** `Random Input Testing results.xlsx` **S-6**

**References** **S-6**

#### 1 Linear Programming Problem Formulation

As discussed, our LP-based algorithm uses a modified version of the Capacitiated Vehicle Routing Problem (CVRP).

Briefly, the objective of the CVRP is to deliver the goods from the depot to all customers using a given number of vehicles that carry the goods, so as to minimise the total cost of all roads that are travelled by the vehicles. Its definition in terms of graph theory is as follows. In a 'road network' graph $G = (V, E)$, the nodes $\{v_1, v_2 \ldots v_n\} \in V$ represent the customers and the edges $\{e_{ij} \mid i, j < n\} \in E$ ($e_{ij}$ is the edge from node $v_i$ to node $v_j$) are the roads, where $cost(e_{ij}) = c_{ij}$ is the cost of travelling the road from the customer $v_i$ to the customer $v_j$. A fleet of $K$ cars is disposable, each of which must start and terminate its journey at the depot $v_0 \in V$ and can serve at most $\kappa$ customers due to vehicle capacity limitations.

The *vehicle flow model* of the CVRP equates determining the vehicles' routes to finding a set of variables $\{x_{ij} \mid 0 \leq i, j \leq n\}$ that indicate which edges have been traversed by some vehicle: $x_{ij} = 1$ if $e_{ij}$ belongs to any of the vehicles' routes and $x_{ij} = 0$ otherwise. Therefore, by introducing dummy variables $u_1, u_2 \ldots u_n$, we can put down a strict LP formulation of the CVRP (S1):

$$\text{minimise } C = \sum_i \sum_j x_{ij} c_{ij}$$

by determining $\{x_{ij}\}$ and $\{u_i\}$ such that

$$\sum_{i=1}^{n} x_{ij} = 1, \quad j = 1 \ldots n \qquad \text{(Constraint 1.1a)}$$

$$\sum_{j=1}^{n} x_{ij} = 1 \quad i = 1 \ldots n \qquad \text{(Constraint 1.1b)}$$

$$\sum_{i=1}^{n} x_{0i} = K \qquad \text{(Constraint 1.2a)}$$

$$\sum_{i=1}^{n} x_{i0} = K \qquad \text{(Constraint 1.2b)}$$

$$u_j - u_i \geq 1 - \kappa(1 - x_{ij}) \; 1 \leq i, j \leq n, \; i \neq j \qquad \text{(Constraint 1.3a)}$$

$$0 \leq u_i \leq \kappa - 1 \;\; 1 \leq i \leq n \qquad \text{(Constraint 1.3b)}$$

Here, Constraints 1.1a-1.1b ensure that every customer is visited by exactly one vehicle (as one incoming and one outgoing edges are selected in any vehicle's route). Constraints 1.2a-1.2b specify that exactly $K$ vehicles leave and return to the depot. Constraints 1.3a-1.3b, using auxiliary variables $u_1 \ldots u_n$, account for both connectivity of the vehicle routes and the vehicle capacity requirements (S1).

Meanwhile, for our pipette tip-saving problem we have a similar graph $G_h = (V, E)$ describing the wells to which the DNA part number $h$ in the sequence of parts to be delivered. The 'goods' are the DNA part solution, the 'customers' are the construct wells (nodes $v_1, v2 \ldots v_n \in V$), each 'vehicle' is a single pipette tip and $cost(e_{ij}) = c_{ij} = 0$ if the same pipette tip can proceed from the well $v_i$ to the well $v_j$ without any cross-contamination. The objective is to deliver a DNA part to all customers requiring it by using as few vehicles as possible. Therefore, the total cost of the vehicle routes is constant at $C = 0$ (including a cost-one edge into any tip's route implies a tip change in the middles of the way, which is a contradiction), while the objective is to minimise $K$, the number of vehicles. This 'swap' of the Objective and Constraint 2 yields the following formulation:

$$\text{minimise } K = \sum_i x_{0i}$$

by determining $\{x_{ij}\}$ and $\{u_i\}$ such that

$$\sum_i x_{ij} = 1, \quad j = 1...n \qquad \text{(Constraint 2.1a)}$$

$$\sum_j x_{ij} = 1, \quad i = 1...n \qquad \text{(Constraint 2.1b)}$$

$$\sum_i \sum_j x_{ij} c_{ij} = C = 0 \qquad \text{(Constraint 2.2)}$$

$$u_j - u_i \geq 1 - \kappa(1 - x_{ij}) \;\; 1 \leq i, j \leq n, \; i \neq j \qquad \text{(Constraint 2.3a)}$$

$$0 \leq u_i \leq \kappa - 1 \;\; 1 \leq i \leq n \qquad \text{(Constraint 2.3b)}$$

The obtained variables $x_{ij}$ allow to uniquely determine every pipette tip's journey. The start of a single tip's route is marked by a variable $x_{0k_1} = 1$. Following the edge $e_{0k_1}$ to the node $v_{k_1}$, we have only one non-zero variable $x_{k_1k_2} = 1$ among those that stand for the edges leaving the node due to Constraint 1b, so the next node in the path is $v_{k_2}$. Similarly, at $v_{k_2}$ the next node is given by the single non-zero variable $x_{k_2k_3} = 1$, and so on until for $v_{k_m}$ we have $x_{k_m0} = 1$. Knowing the order of visiting the wells by a tip, one could easily obtain the sequence of pipette commands to first collect a fresh pipette tip and the relevant DNA part's solution, then deliver aliquots of it to the specified construct wells and finally discard it. Together, these command sequences for every pipette tip, yield the optimised lab robot program to deliver the part number $h$ to all construct wells that need it.

### 2 Determination of the order of DNA part distribution

As every DNA part is assigned its own LP problem, solving which can save pipette tips, the decision which DNA part is to be distributed (to all wells that require it) first, which part the second and so on, is not subject to optimisation by the LP solver. Instead, it is determined by an ordered list $\lambda$ that includes all of the DNA parts.

How can the order of DNA parts be determined in $\lambda$? During algorithm testing, the sequence of DNA parts was random, or alternatively, the parts were given in the order they were encountered in the input. While the former strategy is easy to understand, let us strictly define the latter.

The input to our program pipette implementing the tip-optimising algorithm is a tuple

$\Omega$, where each entry $\omega_i$ (the index signifies that it is found in the the $i$th position in the tuple) is the list of DNA parts comprising a single DNA construct to be assembled – $p_j$ is the $j$th DNA part in this list. Therefore, $\lambda$ is constructed by 'reading through' $\Omega$, as given by Algorithm 1: first, it records into $\lambda$ all parts found in $\omega_1 \in \Omega$; then, it considers $\omega_2 \in \Omega$ and records into $\lambda$ all parts found in $\omega_2$ that have not been recorded in $\lambda$ already; then the program proceeds to record all previously unrecorded parts found in $\omega_3$ and so on until the end of $\Omega$ is reached.

**Algorithm 1** Reading $\lambda$ from $\Omega$

```
λ = ( )
for ω_i in Ω do
    for p_j in ω_i do
        if p_j ∉ λ then
            append p_j to λ
```

As mentioned in Discusson, optimisation of the order of parts in $\lambda$ may further reduce pipette tip consumption in addition to the optimisation provided by our algorithm, which opens up a promising venue for improving the algorithm. As it is impossible to determine the pipette tip savings for every possible $\lambda$ without explicitly running the algorithm with it, such reorderings of $\lambda$ are likely to be heuristic, rather than strictly optimal.

### 3 Annotation to `Start-Stop_Assembly_Random_Inputs.csv`

As Discussed in Results and Methods, for each input size, from just 2 constructs to 96 different construct wells (i.e. the whole well plate), 50 Start-Stop inputs were created, randomly picking one of 6 possible promoters, one of 6 RBSs, one of 3 CDS variants and one of 4 terminators.

In the abstract notation used by the algorithms, a DNA part is given by a pair $(i, j)$, where $i$ is the DNA part type (promoter, CDS, etc.) and $j$ specifies which species of this type it is – in our case, $i = 1$ stood for the promoter parts, $i = 2$ for RBS, $i = 3$ for CDS and $i = 4$ for the terminator. Therefore, each construct well in the file is a single line that has 4 entries, each of which is a pair of numbers encoding the part and its type.

Each input is terminated by a line saying 'end of input' to separate it from the beginning of the next construct well plate description. The inputs for different numbers of wells are separated by a blank line.

This file can also be found in the GitHub repository that contains our algorithm's Python implementation (S2), so that the results presented in the paper could be reproduced.

#### 4 Annotation to `Random Input Testing results.xlsx`

The optimisation of a single input by an algorithm produced a single number, which describes how many pipette tips are required to performed the DNA assembly in an optimised case.

Means, medians and standard deviations of these values were taken across the 50 inputs for every number of wells considered (2 to 96) – the results are displayed in separate tabs. The next three tabs also contain means, medians and standard deviations, but of the percentages of tips saved by the algorithms, which are calculated from the tip consumption data as described in the Methods section.

#### References


(S1) Toth, P., and Vigo, D. *The Vehicle routing problem*; SIAM monographs on discrete mathematics and applications ; no. 9; Society for Industrial and Applied Mathematics: Philadelphia, Pa., 2002.

(S2) Sechkar, K., and Tuza, Z. A. pipette_opt - algorithmic optimisation of an automated liquid-handler's pipette tip consumptions during parallel DNA assembly setup. 2021; `https://github.com/KSechkar/pipette_opt`.
